# Protein engineering, production, reconstitution in lipid nanoparticles, and initial characterization of the *Mycobacterium tuberculosis* EfpA drug exporter

**DOI:** 10.1101/2023.06.26.546575

**Authors:** Olamide Ishola, Adeyemi Ogunbowale, Md Majharul Islam, Elaheh Hadadianpour, Saman Majeed, Oluwatosin Adetuyi, Elka R. Georgieva

## Abstract

*Mycobacterium tuberculosis* (*Mtb*) drug exporters contribute an efficient mechanism for drug resistance. Therefore, understanding the structure–function relationship in these proteins is important. We focused on the *Mtb* EfpA efflux pump, which belongs to the major facilitator superfamily (MSF) and transports anti-tuberculosis drugs outside the bacterial cell. Here, we report on our advancements in producing and characterization of this protein. We engineered a construct of apolipoprotein A-I (apoAI) fused to the N-terminus of EfpA (apoAI-EfpA) and cloned it in an *E. coli* expression vector. This fusion construct was found in a membrane-bound form, unlike the deposited in inclusion bodies EfpA without apoAI. We purified the apoAI-EfpA in detergent to a sufficient degree and reconstituted it in DOPC/DOPS lipids. We found that upon reconstitution in lipid, the apoAI-EfpA forms discoidal protein-lipid nanostructures with a diameter of about 20 nm, resembling nanodiscs. We further detected apoAI-EfpA oligomers in β-DDM and lipid. To the best of our knowledge, this is the first complete protocol on the expression, purification, and lipid reconstitution of the *Mtb* EfpA transported. AlphaFold2 also predicted EfpA oligomers and further bioinformatic analysis confirmed the earlier proposed 14-transmembrane helices of the *Mtb* EfpA. We also found very high identity, >80%, among the EfpA-s of diverse *mycobacterial* species. Outside of *mycobacteria*, EfpA has no close homologues with only low identity with the QacA family of transporters. These findings possibly indicate high specificity of EfpA mechanisms. Our developments provide a foundation for more comprehensive in vitro studies on the EfpA exporter.

## INTRODUCTION

*Mycobacterium tuberculosis* (*Mtb*) is the causative agent of tuberculosis (TB), one of the most devastating and hard-to-control infectious diseases, which leads to severe health complications and often death. Worrisomely, in the last years, the number of drug resistant TB (DR-TB) strains has increased significantly; annually, over half a million new cases of multidrug-resistant TB (MDR-TB) emerge, many of them either extensively or totally drug resistant, rendering them nontreatable with first- and second-line anti-TB antibiotics.^1, 2^ Mutations in the *Mtb*-drug-target genes have been considered a mainstream mechanism to develop DR-TB and MDR-TB.^3-5^ In addition, strong evidence points to *Mtb* membrane exporters’ critical roles in DR-TB and MDR-TB, and strategies to develop inhibitors of these proteins are in progress.^4, 6-10^ In regard to this, the membrane efflux system’s important role in DR and MDR of other pathogenic bacteria is now well recognized, necessitating a comprehensive understanding of how these proteins function to inform on inhibitors’ development.^11-16^

We focused on the *Mtb* drug exporter EfpA (efflux protein A).^17^ EpfA belongs to the major facilitator superfamily (MFS) of transporters, and it expels drugs from the cell for the antiport of H^+^. More narrowly, EfpA belongs to the subfamily of QacA transporters,^17^ which are linked to drug resistance.^18^ Researchers have confirmed EfpA’s role in anti-TB drug export and DR/MDR in several studies: Substantially increased levels (up to 4 times) of EfpA expression in MDR-*Mtb* isolates^7^ and genetically modified *Mtb* strains^19^ upon isoniazid, rifampicin (first-line anti-TB drugs), and ethionamide (second-line anti-TB drug) treatment were detected. EfpA was linked to increased resistance to a combination of isoniazid and ethambutol.^20^ It was found further that the overexpression of EfpA protein from *M. smegmatis*, which shares 80% identity with the *Mtb* EfpA, induces high drug tolerance (MDR) to several first- and second-line anti-TB drugs, including rifampicin (observed 4-fold increased resistance), isoniazid, streptomycin, amikacin (16-fold increase), and ethidium bromide.^21^ Therefore, the role of *Mtb* EfpA in DR and MDR is evident. This protein is now recognized as a very specific *mycobacterial* druggable target, prompting the search for efficient EfpA inhibitors.^22^ To this end, comprehensive studies to understand EfpA-anti-TB drug interactions and transport will be very informative. However, such studies have been very limited, possibly because of the unavailability of recombinantly expressed and purified EfpA.

Here, we report our efforts to overcome the deficiencies in the *in vitro* studies of *Mtb* EfpA by developing strategies to express this protein in *E. coli* and purify it sufficiently for further studies. To do so, we utilized protein engineering to generate a chimera construct containing the apolipoprotein A-I (apoAI) fused to the N-terminal of EfpA (apoAI-EfpA construct), which enabled us to express the protein in an *E. coli* membrane-bound form. To the best of our knowledge, this is the first complete protocol for the heterologous expression and purification of *Mtb* EfpA reported. Interestingly, when the apoAI-EfpA was reconstituted in lipid membranes comprising DOPC (1,2-Dioleoyl-sn-glycero-3-phosphocholine) and DOPS (1,2-dioleoyl-sn-glycero-3-phospho-L-serine), we found by negative staining electron microscopy (nsEM) that it forms protein-lipid discoidal nanoparticles resembling nanodiscs, presumably due to the apoAI moiety.^23, 24^ To enhance our studies on *Mtb* EfpA, we used bioinformatic tools to predict EfpA membrane topology, confirming its 14-transmembrane helices’ organization.^25^ We also studied the degree of conservation of EfpA exporters among *mycobacterial* species and found a high degree of identity (greater than 85% among the studied species), but we found only about 20% identity between Mtb EfpA and homologues of the QacA subfamily. This finding again confirms these proteins’ exclusive specificity to *mycobacteria*. The results from size exclusion chromatography (SEC) and negative staining electron microscopy (nsEM) indicated possible protein oligomerization, which was supported by further assessment using the AlphaFold2 molecular modeling software for EfpA in inward facing conformation. SWISS MODEL software generated structural models of the monomeric *Mtb* EfpA in both inward and outward facing conformation, using available high-resolution structures of distant protein homologues.

## RESULTS

### The expressed in *E. coli* chimera construct apoAI-EfpA is a membrane-bound protein

In our initial attempts to produce in *E. coli* the full-length (FL) EfpA with just a polyhistidine (His_8_) tag as well as with a combined His_8_ and FLAG tag, we found that upon expression, the protein was deposited in the insoluble inclusion bodies, similar to other *Mtb* membrane proteins.^26^ Therefore, the further purification of this protein required inclusion bodies’ solubilization using high concentration of sodium dodecyl sulfate (SDS) and protein refolding by stepwise addition of milder detergent n-dodecyl-β-D-maltoside (β-DDM) while removing SDS (Supporting Information and Supporting Figure 1).

**Figure 1.**
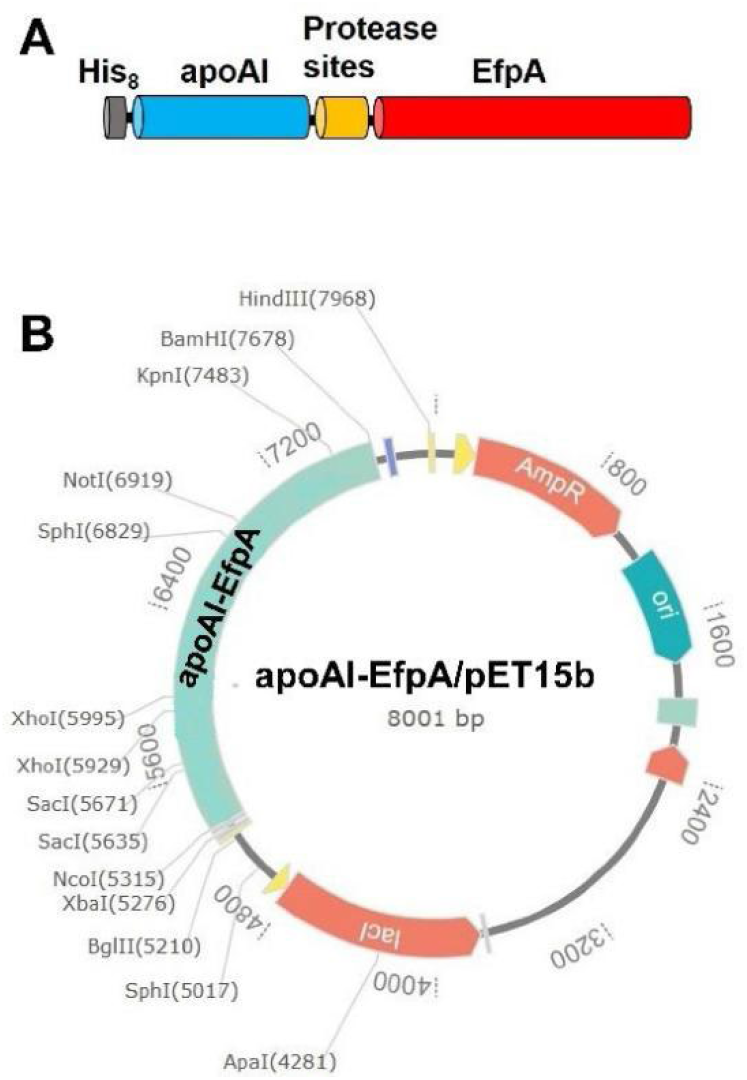
Design and cloning of the apoAI-EfpA chimera construct. (A) A cartoon representation of the apoAI-EfpA construct: His8+apoAI tag (in gray and blue, respectively) was fused to the N-terminus of EfpA; the protease recognition sites between apoAI and EfpA are in orange; EfpA is in red. (B) The gene encoding the apoAI-EfpA was cloned in pET15b vector between the NcoI and BamHI restriction sites resulting in an apoAI-EfpA/pET15b protein expression vector. The vector map was created using the GenSmartTM (GenScript) software.

This requirement prompted us to look for alternative strategies to produce EfpA in a more native state. To do so, we engineered a chimera construct of apolipoprotein A-I (ApoAI), relying on previously developed protocols using apoAI fused to the C-terminus of several integral membrane proteins to produce them in a soluble form.^27^ However, we fused the apoAI to the N-terminus of EfpA (apoAI-EfpA construct) (Figure 1A and Supporting Figure 2). In the linker between apoAI and EfpA, we engineered two recognition sites for SUMO and Thrombin (Thr) proteases. This construct’s total calculated molecular weight is 84.8 kDa. We cloned the gene encoding this chimera construct in the pET15b expression vector (Figure 1B). When it is expressed in *E. coli*, we expected to find the protein in the soluble cytoplasmic fraction, as in the previous study.^27^ Surprisingly, when we screened all the cell fractions (i.e., cytoplasmic, membrane, and inclusion bodies), the membrane fraction contained most of the protein (Supporting Figure 2). Therefore, we proceeded with handling the apoAI-EfpA as a membrane protein.

**Figure 2.**
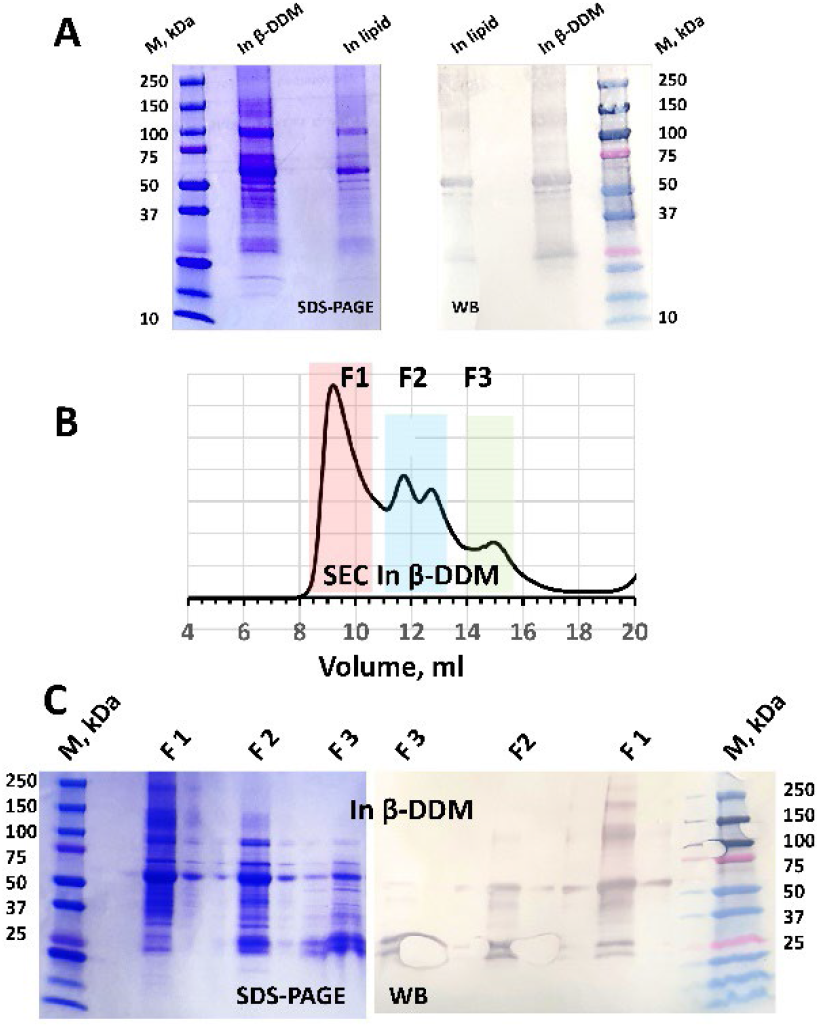
Assessment of the purified apoAI-EfpA. (A) SDS-PAGE and WB of Ni-+Co-affinity purified protein in β-DDM and in lipid are shown. (B) SEC of the protein in (A)—three main fractions F1, F2 and F3 highlighted in red, blue, and green, respectively, were collected. (C) SDS-PAGE and WB of the protein in F1, F2 and F3 in (B). The additional bands seen between the wells of F1, F2 and F3 on the SDS-PAGE and WB are due to leaky gels.

Therefore, we solubilized the membranes by using high concentration (30-35 mM) of β-DDM and purified the apoAI-EfpA using Ni-affinity and Co-affinity (Ni-+Co-affinity) chromatography sequentially. The isolated protein was of sufficient purity (Figure 2). In addition to the bands at the molecular weight of apoAI-EfpA, SDS polyacrylamide gel electrophoresis (SDS-PAGE) and western blotting (WB) displayed His-tag–positive protein bands at molecular weight below that of the apoAI-EfpA monomer, which could be a result of fractional protein degradation but could also be due to anomalous migration of a more folded apoAL-EfpA in SDS with an effectively smaller protein molecular weight. The latter is plausible because we observed similar low-molecular-weight protein bands for the ApoAI-free EfpA constructs purified from inclusion bodies (Supporting Figure 1). What is more, SDS-PAGE- and WB-positive protein bands at molecular weight of about 150 kDa (close to the molecular weight of an apoAI-EfpA dimer) were also observed (Figure 2A,C), which points to apoAI-EfpA oligomerization.

Still, to ensure the high protein homogeneity, we subjected the Ni-+Co-affinity purified apoAI-EfpA to SEC and found the protein mostly in two fractions—fractions 1 and 2 (F1 and F2) in Figure 2B. Of these, the SEC peak of F1 was well-defined and corresponds to protein of molecular weight greater than 200 kDa and F2 was a combination of two peaks containing proteins of molecular weight greater than 100-150 kDa.^28^ Both F1 and F2 were WB-positive, thus containing apoAI-EfpA. This result provides one more evidence of apoAI-EfpA’s homo-oligomerization in detergent, but these oligomers a somewhat heterogeneous ranging from tetramers to lower order oligomers. It is worth noting that the F1 peak occurs well after the void volume of 6 ml for the column we used for SEC.^28^ Similarly to Ni+Co-affinty purified protein, the SDS-PAGE and WB of the F1 and F2 (Figure 2C), showed protein bands at molecular weight of monomeric and oligomeric FL AI-EfpA. However, WB-positive protein bands corresponding to molecular weight lower than that of the FL apoAI-EfpA ware also observed. Here we rationalize that if these bands correspond to truncated apoAI-EfpA, it is unclear to us whether the proteolysis takes place before or during the SDS-PAGE procedure, given that the SEC peak of F1 is well-shaped and indicates high protein homogeneity. As discussed above, anomalous protein migration is also possible.

### Negative-staining electron microscopy suggests that apoAI-EfpA forms a homo-oligomer, and when reconstituted in lipid assembles into nanodisc-like protein-lipid particles

We subjected the Ni+Co-affinity purified apoAI-EfpA in β-DDM to nsEM and obtained clear images of protein particles, which are apparently larger than expected for a protein of about 84 kDa (Figure 3A, Supporting Figure 3). This result prompted us to explore AI-EfpA’s ability to form oligomers. To gain further insights into the oligomeric state of apoAI-EfpA, we labeled the N-terminus His-tag of the protein with 5 nm gold nanoparticles (GNP-s). The nsEM images of this sample clearly show clusters of protein-associated GNP-s ranging in number from 1 to 4 (Figures 3B and C), suggesting that apoAI-EfpA is indeed oligomeric. Interestingly, we did not observe protein clusters with more than 4 GNPs, which might indicate that the tetramer is the highest-order oligomer of this protein in β-DDM. One consideration here is that the concentration of GNPs in the studied samples was significantly smaller than the protein concentration (18 µM protein/0.14 µM GNP). Therefore, it is reasonable to expect that most of the protein molecules do not have a GNP attached to them. If further consider statistical distribution of protein-bound GNP-s in the apoAI-EfpA oligomers, observing oligomers with 1, 2, or 3 protomers having GNP-s as well as protein oligomers having no GNP-s is also expected.

**Figure 3.**
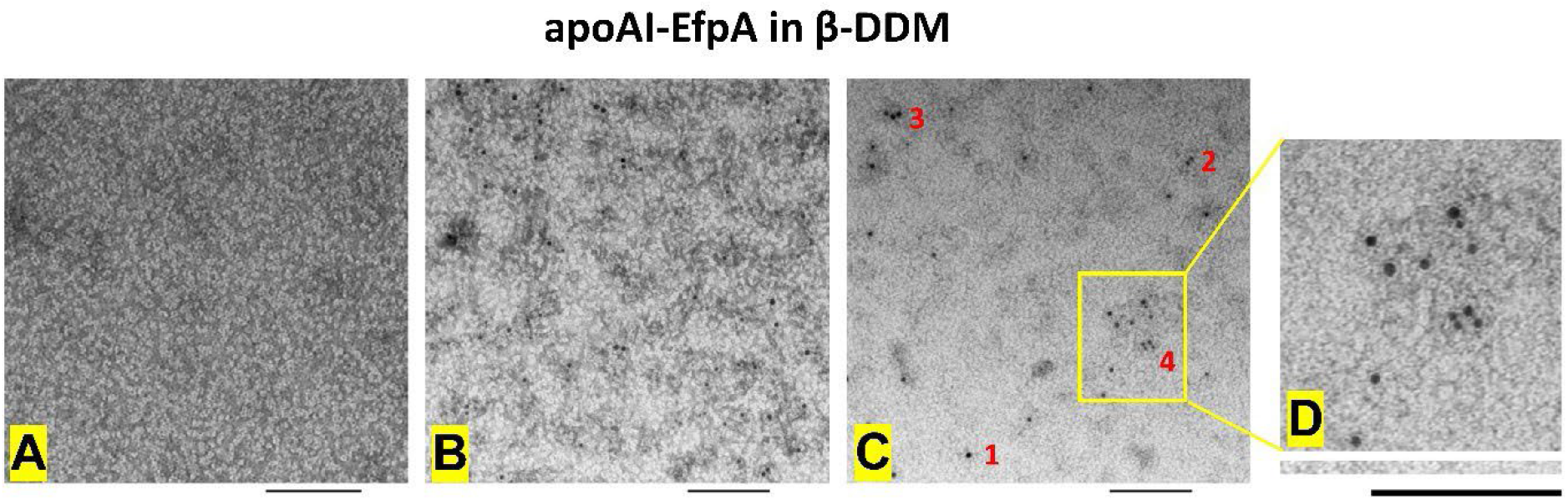
nsEM images of apoAI-EfpA in β-DDM. (A) A selected image of a protein sample without GNPs; (B) and (C) Selected images of protein labeled with GNP-s (black dots); (D) An expanded view of the image region enclosed in the yellow rectangle in (C). The scale bar in all images is 100 nm. In all images, apoAI-EfpA oligomers are visible, particularly exemplified by the clusters of protein-bound GNP-s. The red numbers in (C) show the apparent number of GNP-s within a protein cluster. Note, the number of GNP-s does not report directly on the number of protein monomers, as the concentration of GNP-s in the imaged sample was about 130 times less than the concentration of protein. Still, the grates number of GNP-s observed is 4, which might indicate that this is the highest order of the apoAI-EfpA oligomer.

We further reconstituted the apoAI-EfpA in a mixture of DOPC/DOPS (a 70:30 molar ratio). To ensure the protein incorporation in the lipid environment, we first solubilized the lipid bilayers (multilamellar vesicles) with Triton-100 and then added to them the apoAI-EfpA in β-DDM. After incubating this mixture for 1 h at 22 °C, we gradually removed the detergents.^29, 30^

Using nsEM, we again visualized the proteolipid samples with and without GNP-s attached to the protein. In this case, we observed discoidal protein-lipid nanoparticles with an average diameter of about 20 nm (Figure 4 and Supporting Figure 3). These structures resemble proteins incorporated in nanodiscs, which are also lipid nanoparticles stabilized by a belt of a scaffold proteins (apolipoprotein AI).^31-33^ Therefore, because our chimera construct contains apoAI, it is not surprising that nanodisc-like particles were formed. Still, in the conventional case, 2 copies of the scaffold proteins are necessary to stabilize a single nanodisc,^31^ but for the apoAI-EfpA, we observed up to 4 protein-bound GNP-s in a single discoidal structure (Figures 4C, D, and E), suggesting a tetrameric protein. Similar to apoAI-EfpA in β-DDM, under our experimental conditions, the GNP-s concentration was substantially lower than the protein concentration. Therefore, not all apoAI-EfpA monomers have a GNP attached to them.

**Figure 4.**
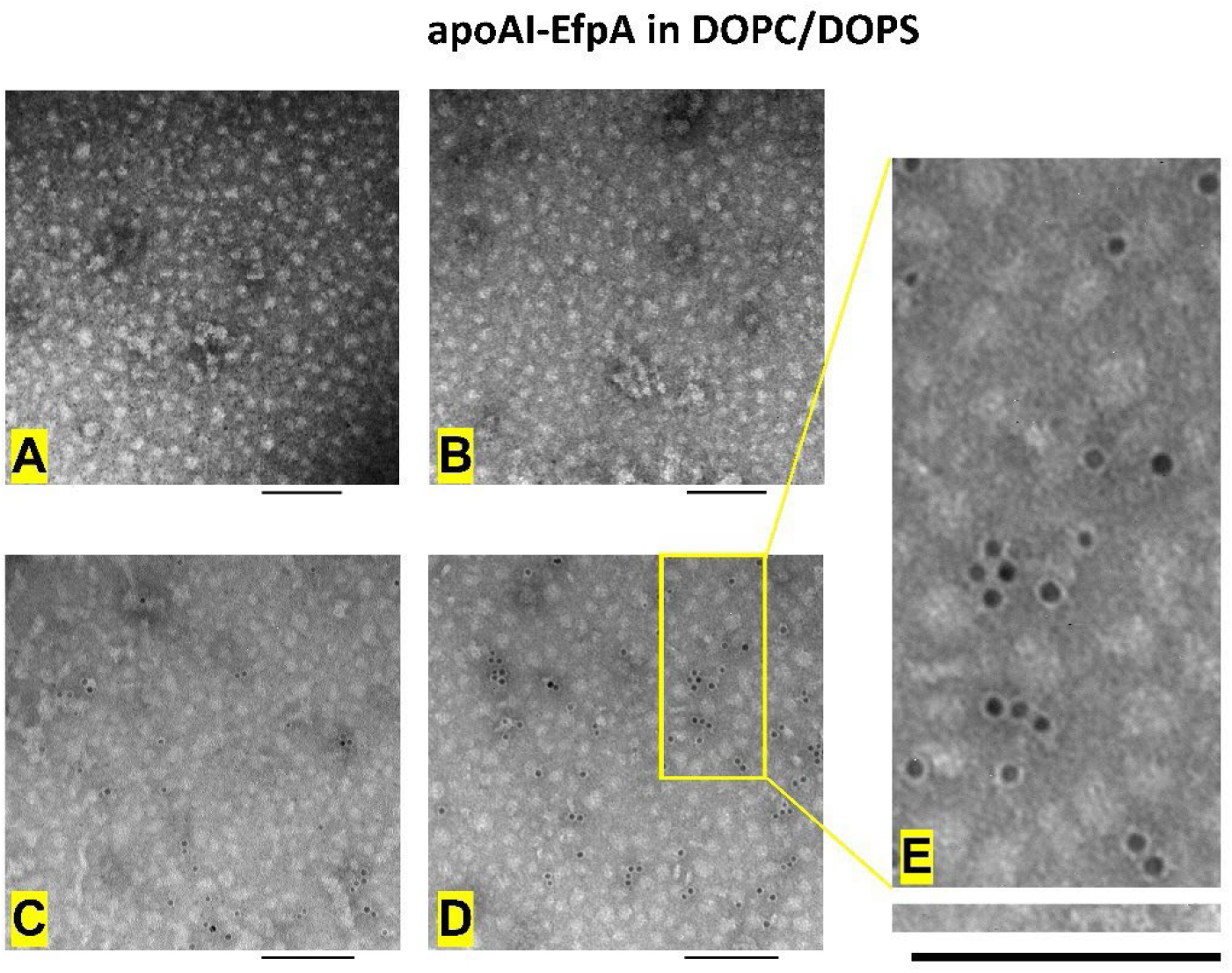
nsEM images of apoAI-EfpA in lipid. (A) and (B) Selected images for protein with no GNPs; (C) and (D) Selected images for protein labeled with GNPs (black dots); (E) An expanded view of the image region enclosed in the yellow rectangle in (D). The scale bar is 100 nm. All images show that upon reconstitution in DOPC/DOPS lipids, apoAI-EfpA formed nano-sized (about 20 nm in diameter) protein-lipid particles. Again, protein oligomers with order of up to four (based on the number of clustered GNPs) was observed.

We further reconstituted the apoAI-EfpA from SEC F1 (Figure 2) in DOPC/DOPS, under the same condition as the reconstitution of the Ni-+Co-affinity purified protein. Again, the protein-lipid discoidal structures with varying AI-EfpA homo-oligomers were observed again (Figure 5).

**Figure 5.**
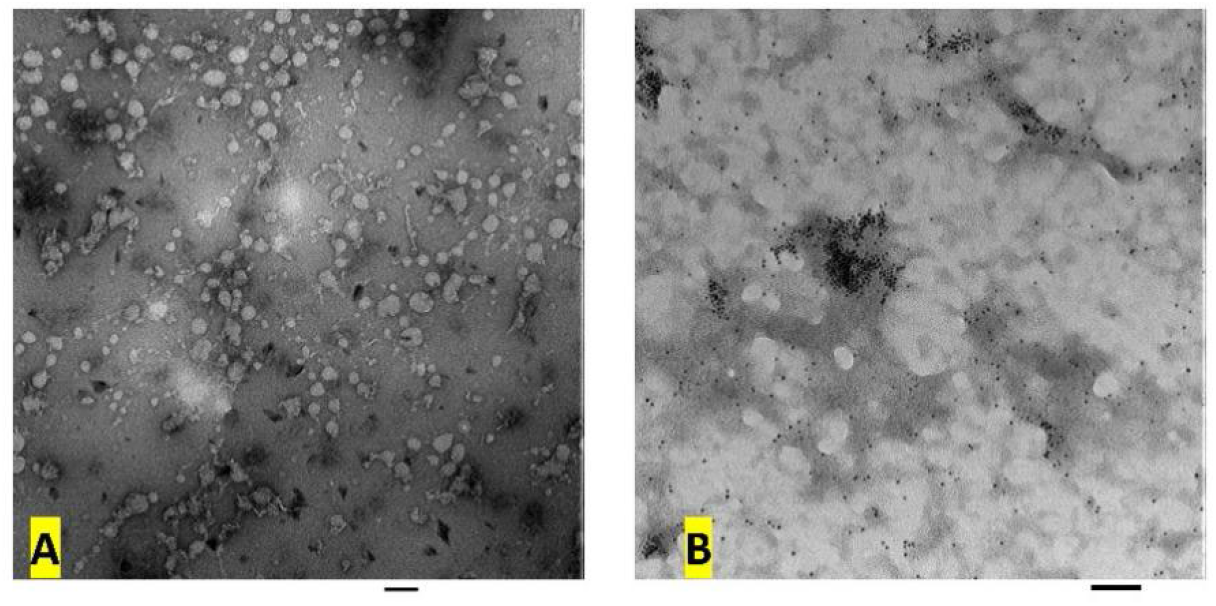
nsEM images of apoAI-EfpA from SEC F1 in lipid. (A) and (B) Selected images for protein without and with GNPs, respectively. The scale bar is 100 nm. The images show that upon reconstitution in DOPC/DOPS lipids, apoAI-EfpA formed nano-sized protein-lipid particles. Again, protein oligomers with order of up to four (based on the number of clustered GNPs) was observed. Some of GNPs were not bound to the protein and segregated outside of the lipid in (B).

Besides confirming the tendency of EfpA to self-associete, our observations might suggest that in lipid this protein forms a sufficiently tight oligomer, which is able to restructure the nanodiscs to include up to 4 copies of apoAI (which is covalently linked to EfpA). If this is the case, our results might indicate that nanodiscs could possess adequate structural and size flexibility to accommodate membrane proteins of various sizes. Further studies will provide more detail on this matter. Another important aspect of this observation is that we were successful in creating a chimera protein that carries the transmembrane protein and the scaffold protein, which can form proteolipid nanoparticles on its own. The designed protease sites (i.e., SUMO and Thr) could provide even more flexibility in the system by separating the apoAI tag from EfpA, which will be explored in future studies.

### Bioinformatic analysis confirms that *Mtb* EfpA is a 14-transmembrane-helix exclusively *mycobacterial* MFS transporter, which possibly forms homo-oligomers

To expand our studies and supplement our experimental data, we conducted a bioinformatic analysis of the *Mtb* EfpA protein.

#### The EfpA exporter has a 14-transmembrane-helix topology

To assess the topology of the membrane-residing *Mtb* EfpA exporter, we utilized the *Protter* proteoform visualization program,^34^ which predicted 14-transmebrane helices (TMs) with intracellular location of the N-and C-termini (Figure 6). This is in agreement with the prediction regarding EfpA,^17^ and the terminus locations coincide with those of other MSFs with 14- and 12-TMs.^35-37^

**Figure 6.**
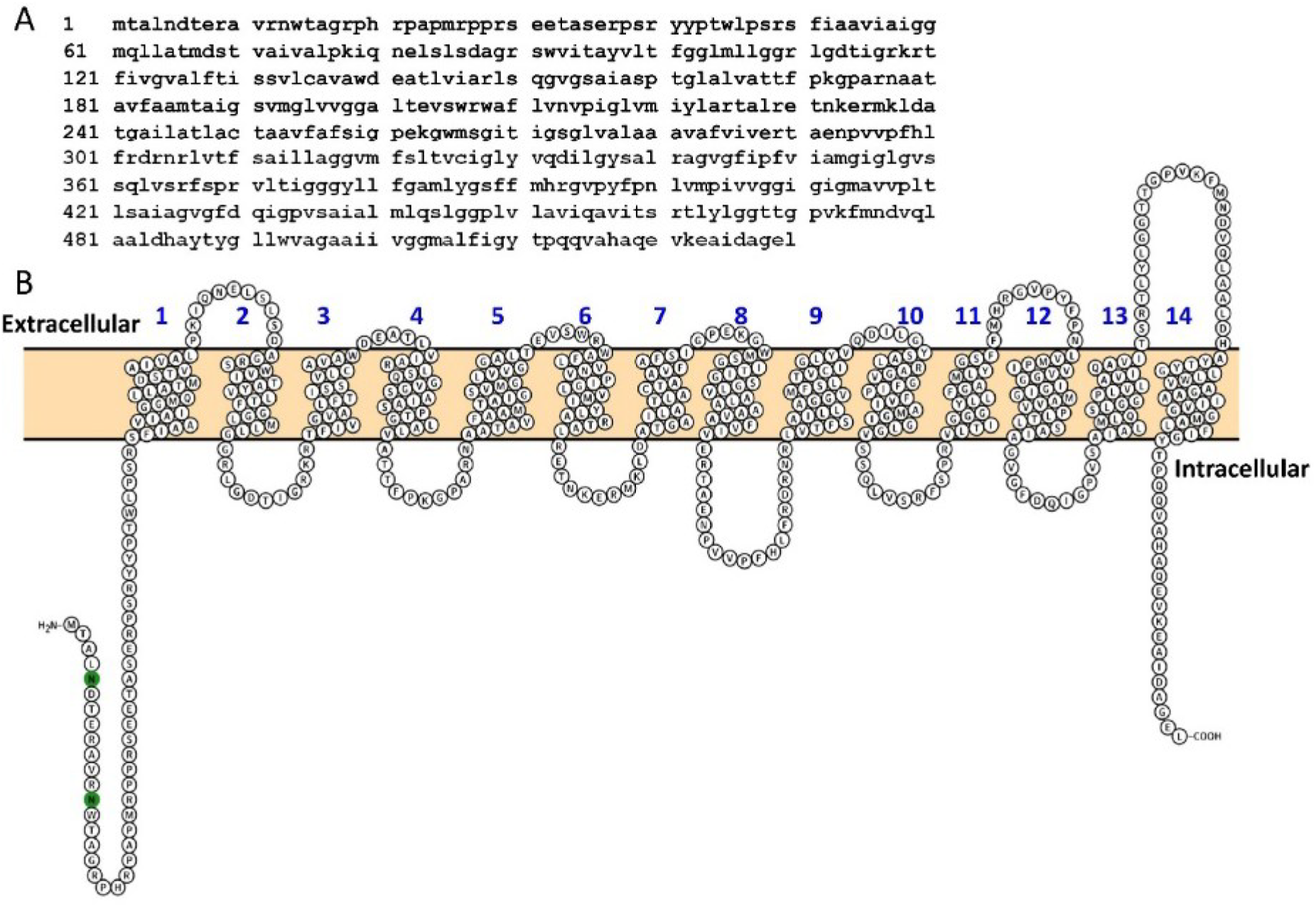
The *Mtb*-encoded EfpA efflux pump. (A) Amino acid sequence and (B) Predicted protein topology—TMs are numbered from 1 to 14 in blue; N- and C-termini have intracellular (cytoplasmic) location.

#### The amino acid sequence of EfpA exporters is highly conserved among mycobacterial species but shares low similarity with the QacA family

Because EfpA is a member of the QacA family^17^ but is mostly limited to *mycobacteria*,^8, 17^ we aimed to determine the degree of identity between *Mtb* EfpA with homologues from other *mycobacterial* species and among the QacA exporters. Based on the amino acid sequences’ alignment, we found that the EfpA efflux pumps encoded by several selected *mycobacterial* species are highly conserved: Among all the *mycobacterial* species we studied, the identity of EfpA proteins is greater than 85%. This result is in agreement with the literature.^17^ Our alignment included EfpA-s from *M. tuberculosis*, *M. shinjukuense* (causes pulmonary infection^38^), *M. simiae* (infection leads to bronchiectasis, micronodular lesions, and weight loss^39^), *M. szulgai* (also causes lung diseases^40^), *M. arosiense*, and *M. intracellulare* (Figure 7). On the other hand, our analysis of the amino acid sequences of selected QacA family proteins (QacA_*S. Aureus*, EmrB/QacA_*B. Pseudomallei*, EmrB/QacA_*C. Disporicum*, etc.) (Figure 8) resulted in relatively low identity, ca. 20%, of *Mtb* EfpA with the rest of the studied proteins. These results, together with the limited occurrence of EfpA in bacterial genomes, emphasize this protein’s exclusiveness to *mycobacteria* and might suggest unique details in its mechanism.

**Figure 7.**
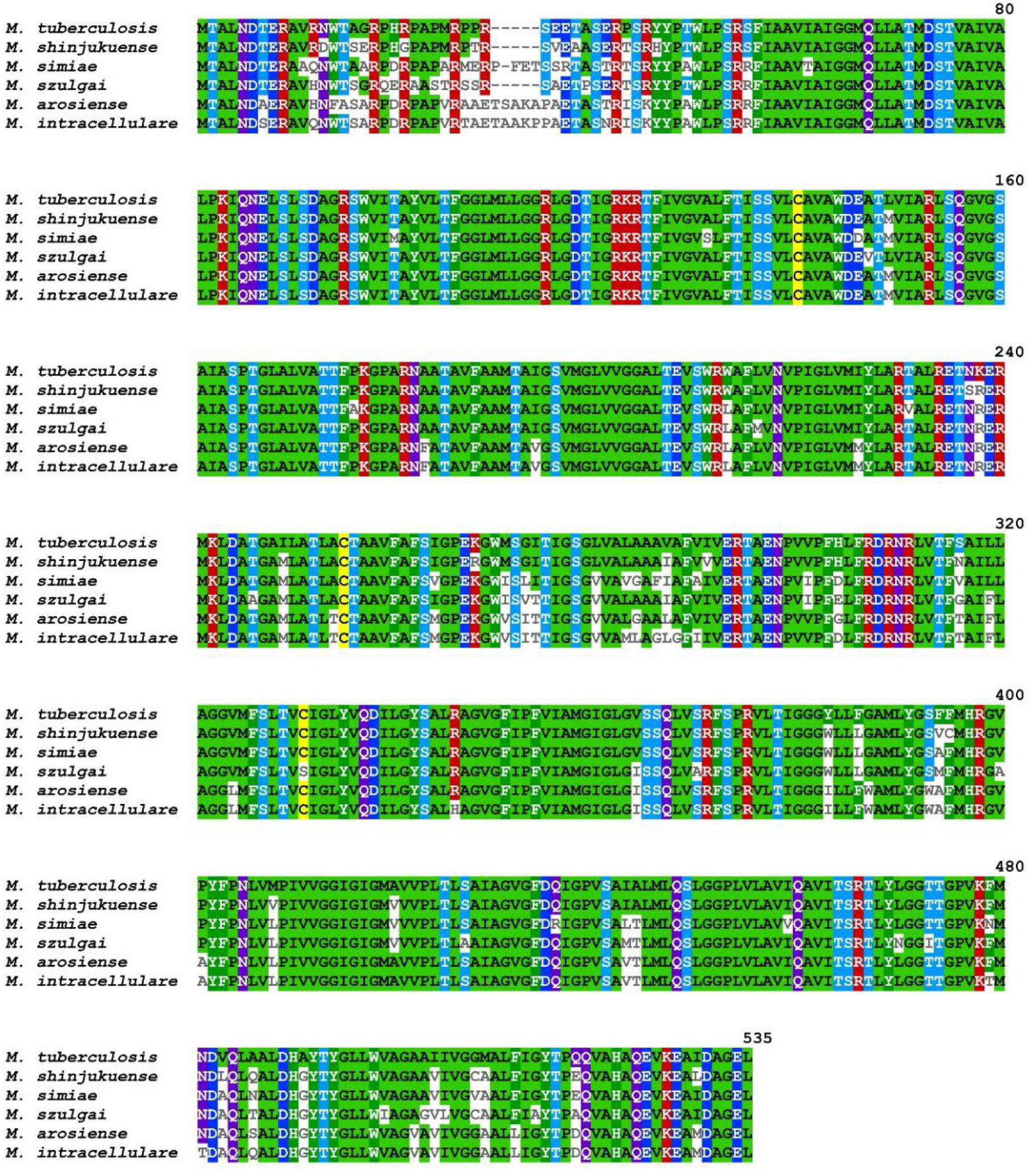
The amino acid sequence alignment of EfpA proteins encoded by selected *mycobacterial* species, which are *M. tuberculosis*, *M. shinjukuense*, *M. simiae*, *M. szulgai*, *M. arosiense*, and *M. intracellulare*. The T-COFFEE Multiple Sequence Alignment software was used to perform the analysis and visualize the data. The amino acids are colored by identities.

**Figure 8.**
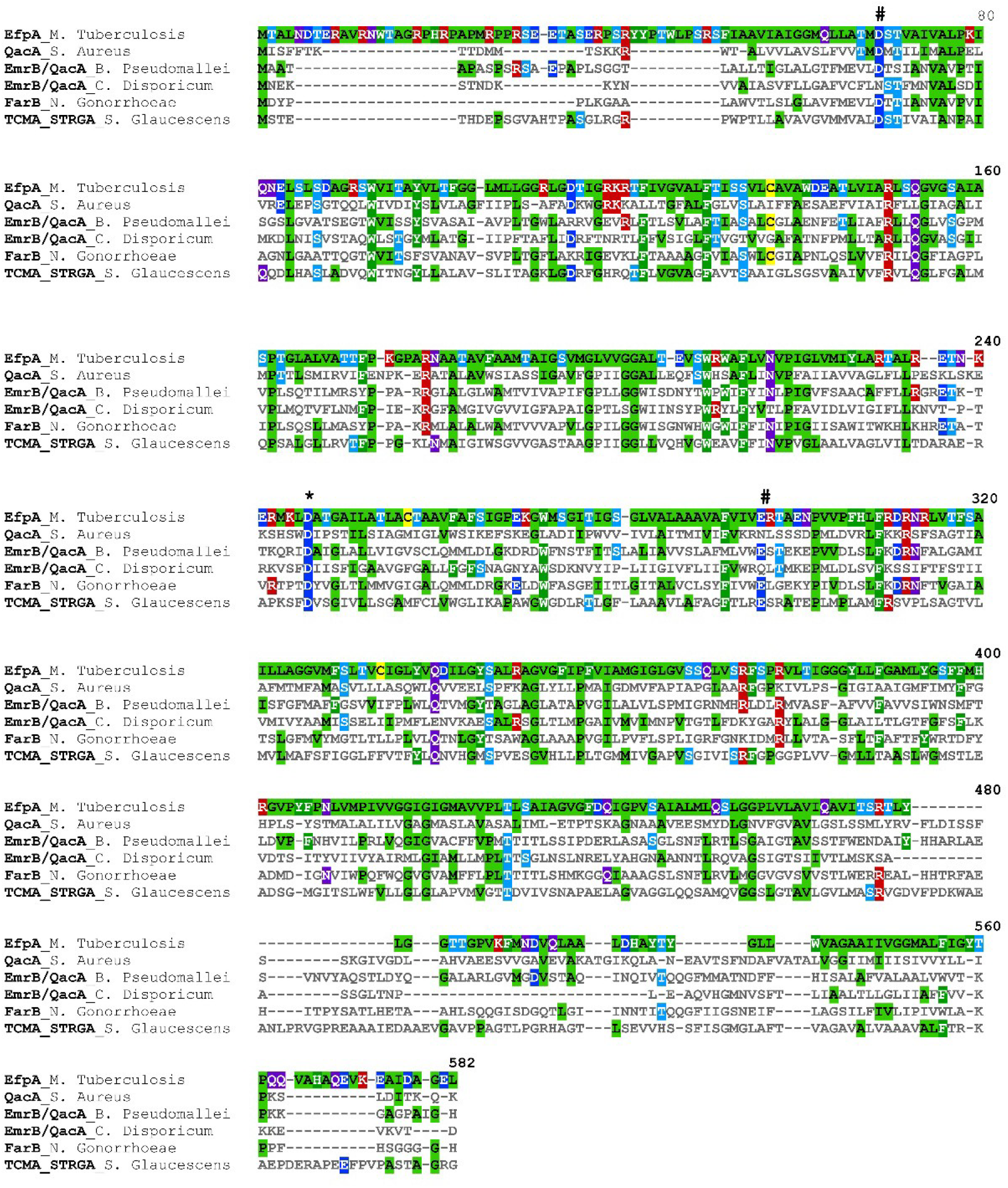
Amino acid sequence alignment of selected members of the QacA family of exporters, which are *Mtb* EfpA, *S. Aureus* QacA, *B. Pseudomallei* EmrB/QacA, *C. Disporicum* EmrB/QacA, *N. Gonorrhoeae* FarB, *S. Glaucescens* TCMA_STRGA. The T-COFFEE Multiple Sequence Alignment software was used to perform the analysis and visualize the data. The amino acids are colored by identities.

#### AlphaFold2-predicted Mtb EfpA dimerization

Because our experimental data suggested possible oligomerization of apoAI-EfpA, which could be contributed by the EfpA moiety, we conducted AlphaFold2 multimer^41^ analysis to possibly predict EfpA self-association based on EfpA’s amino acid sequence and database homology models. Indeed, AlphaFold2 predicted possible EfpA dimer formation (Figure 9A); however, with much higher confidence of the prediction of EfpA monomer in inward-facing conformation vs. EfpA dimer, based on the obtained PAE plot (Figure 9B).

**Figure 9.**
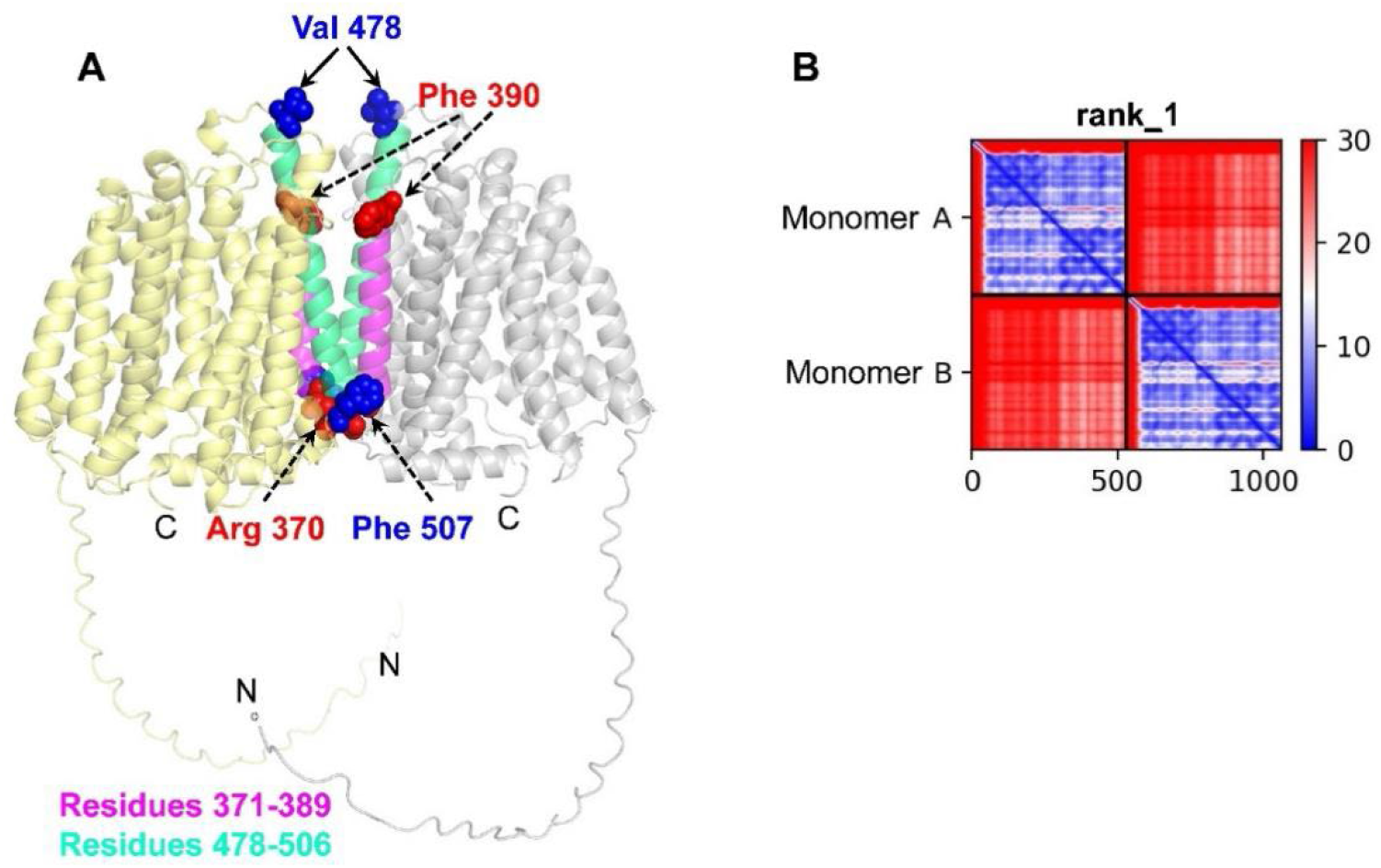
The AlphaFold2-predicted *Mtb* EfpA dimer. (A) Structural model of EfpA dimer— the possible dimerization might occur via contacts in TM helices 11 (residues 371-389) and 14 (residues 478-506). (B) The PAE plot showing level of confidence in relative positions of residues (high is in blue, medium is in white, and low is in red). While the prediction of monomer structure is with relatively high confidence, the prediction of dimerization contacts is highly uncertain with increased average dimerization confidence for the EfpA region after residues 350.

Perhaps because of the limited availability of structural templates in AlphaFold database due to only few solved high-resolution structures of MFS superfamily transporters containing more than 10 TM helices, particularly of those from the QacA sub-family, only the structure of EfpA in inward-facing conformation was predicted. Because of these reasons and due to currently limited applications of AlphaFold2 to transmembrane proteins and their homo-oligomers,^42^ the obtained model should be interpreted with caution. Still, the AlphaFold2 predicted oligomerization is in agreement with our experimental results.

#### SWISS-MODEL predicts structural models of Mtb EfpA in outward- and inward-facing conformations and reveals possible structural rearrangements

We used the SWISS-MODEL program^43^ to generate the 3D structural models of *Mtb* EfpA (Figure 10). As templates, we used the 2 existing X-ray structures of the *S. aureus* NorC (PDB#7D5Q) and *E. coli* DtpA (PDB#6GS1) in outward- and inward-facing conformation, respectively.^36, 37^ The models revealed large conformational changes upon transition from outward- to inward-facing states underlying the binding and translocation of substrate (drug/H^+^). These conformational isomerizations are in agreement with the “rocking-switch” mechanism of the MSF transporters.^35, 44^ The generated models would be useful in further characterization of the *Mtb* EfpA structure–function relationship. However, experimental and detailed computational data would also be needed to elucidate the specificity in the functional mechanism of the *Mtb* EfpA because this protein shares less than 20% and 30% identity with NorC and DtpA, respectively, as our analyses revealed; it is also very specific for *mycobacteria*, as mentioned above.

**Figure 10.**
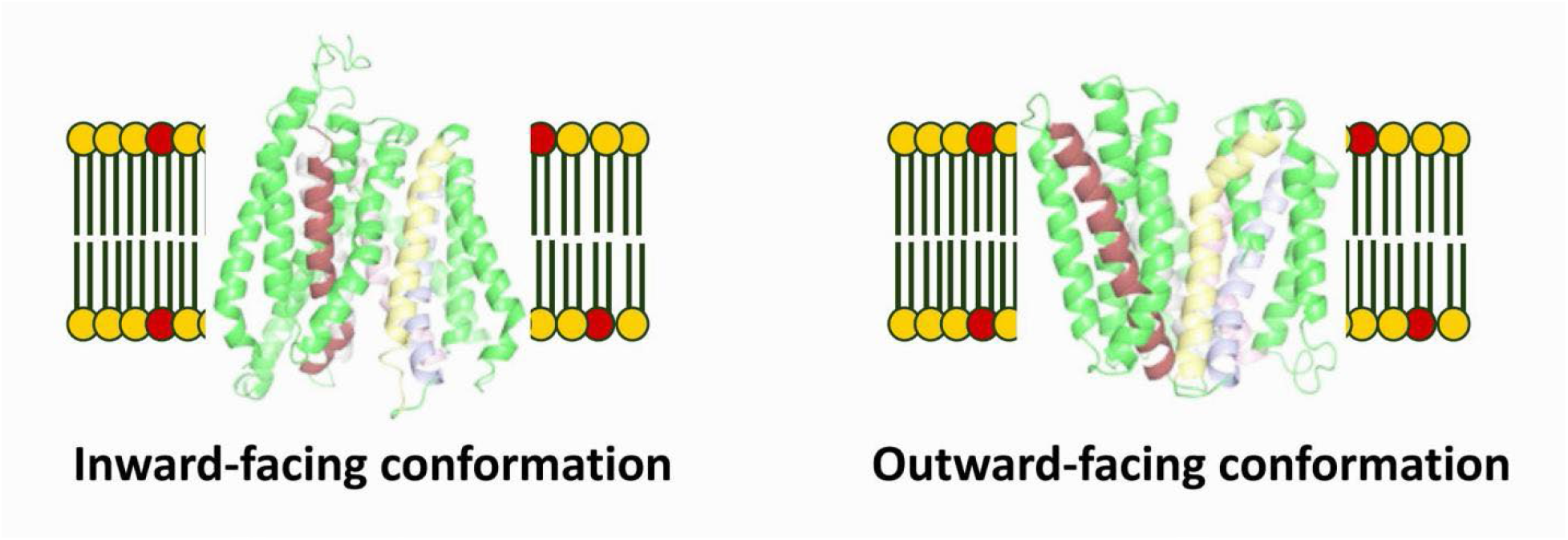
Structural models of *Mtb* EfpA in inward- and outward-facing conformations based on the X-ray structures of NorC (PDB#7D5Q) and DtpA (PDB#6GS1). The TMs undergoing large conformational isomerization upon transition from inward to outward facing conformation based on these models are colored as follows: residues 89-115 are in pink, residues 141-169 are in light-blue, residues 174-202 are in yellow, residues 397-426 are in ruby. Membrane lipid headgroups are in yellow-orange.

## DISCUSSION

Understanding the structure, conformational dynamics, drug specificity, and drug translocation mechanism of *Mtb* drug exporters is important for the development of informed strategies to inhibit these proteins. However, most of these proteins have not been produced in purified form, hindering the in vitro studies. Here, we focus on the *Mtb* EfpA drug exporter. Several drug substrates of EfpA, including first- and second-line anti-TB drugs, were proposed, and the literature provides solid evidence of this protein’s role in DR and MDR development.^7, 19-21^ However, currently, molecular details about the mechanism(s) of EfpA are very limited; no protocols to produce and isolate this protein have been published except our recent short report.^45^ In the current study, we made significant progress by developing strategies to produce this protein in an *E. coli* host recombinantly and purify it to a high degree. To do so, we engineered a chimera construct apoAI-EfpA that had plasma membrane localization when expressed in *E. coli*, which enabled us to handle apoAI-EfpA as a membrane protein (i.e., to extract it from the membrane using detergent, purify it in detergent, and then reconstitute it in lipid membranes). To the best of our knowledge, this is the first protocol reported for the expression and purification of the FL *Mtb* EfpA drug exporter. Such an achievement could be an important stepping stone in the in vitro studies of EfpA because it will open up future opportunities to elucidate the structure–function relationship of this protein. In addition, the apoAI-EfpA construct has the advantage of forming protein-lipid discoidal nanoparticles, similar to the nanodiscs used in membrane protein studies.^23, 24, 46^ This could make it possible to characterize EfpA’s properties in a system comprising just 2 components, the apoAI-EfpA and lipid. The methodology could be further explored in studies of other membrane proteins.

Another interesting observation in our study is the oligomerization of EfpA in detergent and lipid, with tetramer being the largest oligomer visualized by nsEM. This finding might suggest that the *Mtb* EfpA functions as an oligomeric protein, but further details would be necessary. EfpA is a MFS transporter,^6, 17^ and majority of this family members typically function as monomers with no need of oligomerization for substrate translocation.^35, 44^ Still, oligomerization of MFS membrane transporters was observed previously (e.g., the human proton-coupled folate transporter^47^ and the plant nitrate transporter, NRT1.1).^48^ Thus, the oligomerization of EfpA is not completely surprising and it might play a stabilizing or regulatory role, which could be determined in subsequent studies.

## MATERIALS AND METHODS

### Cloning, expression, and purification of *Mtb* EfpA as a fusion construct with apoAI

Three fusion constructs of *Mtb* EfpA were designed: His_8_-linker-*Mtb* EfpA, His_8_-linker-FLAG-linker-*Mtb* EfpA, and His_8_-linker-apolipoprotein A-I (apoAI)-linker-SUMO protease site-linker-Thrombin protease site-linker-*Mtb* EfpA. The DNA encoding each of these constructs was commercially synthesized and cloned in pET15b vector at NcoI and BamHI sites (*GenScript*, Inc) and then transformed into Bl21(DE3) *E. coli* cells. However, upon expression in *E. coli*, we found that the first 2 fusion proteins were deposited in the insoluble inclusion bodies^49^ (Supporting Information). Therefore, we proceeded with the third construct, in which apoAI was fused to the N-terminus of *Mtb* EfpA. Throughout the text in this paper, we refer this construct as apoAI-EfpA. After transforming the plasmid DNA encoding apoAI-EfpA into *E. coli* BL21(DE3) chemically competent cells, colonies were grown on LB/agar/100 µg/ml Ampicillin (Amp) plates overnight at 37 °C. The next day, 200 mL of LB medium supplemented with 100 µg/ml Amp were inoculated with a single colony of transformed cells, and an overnight culture was grown at 37 °C in a shaker incubator. The next day, 30 mL of this overnight culture was transferred into large flasks containing 2 L LB medium supplemented with 100 µg/ml Amp. The cells were further grown at 37 °C in a shaker incubator until the cell culture OD reached 0.6-0.8; then, the temperature was decreased to 14 °C, and the protein expression under the control of the T7 promoter was initiated by adding 0.5 mM IPTG (Isopropyl ß-D-1-thiogalactopyranoside) to the cell culture. The expression proceeded overnight at 14 °C for about 16 h.

The next day, the cells were harvested by centrifugation at 4,100 rpm in an Avanti J-15R centrifuge (Backman Coulter) and resuspended in a buffer containing 20 mM HEPES pH 7.4, 300 mM NaCl and 5% glycerol (Buffer A). All following procedures were carried out either on ice or at 4 °C. PMSF (phenylmethylsulfonyl fluoride), lysozyme, and TCEP (tris(2-carboxyethyl)phosphine) were added to the cell solution to 1 mM, 0.6 mg/ml, and 200 µM final concentrations, respectively. The cells were broken open by sonication. The unbroken cells, cell debris, and inclusion bodies were spun down in an Avanti J-15R (Beckman Coulter) centrifuge at 5,500 rpm for 15 min at 4 °C, and then the cell membranes were spun down in an Optima XE-90 ultracentrifuge (Beckman Coulter) in a rotor type 70.1Ti at 40,000 rpm (146,550.4xg) for 1 h at 4 °C. All fractions (i.e., cell debris with inclusion bodies, membranes, and soluble) were subjected to SDS-PAGE and WB to determine the location of the expressed apoAI-EfpA. In several repeated experiments, we found the protein only in the membrane fraction because the soluble and insoluble fractions were WB-negative for this protein.

Afterward, we followed the procedure for membrane solubilization and protein extraction in detergent n-Dodecyl-beta-Maltoside (β-DDM), as previously described.^29^ The detergent-solubilized apoAI-EfpA was purified using Ni-affinity and Co-affinity chromatography techniques sequentially. Briefly, the protein mixture was incubated with Ni-NTA resing (*Quagene*) for 1 h at 4 °C, the fraction of unbound protein was discarded, and the Ni-NTA resin with bound apoAI-EfpA was washed with 12 resin volumes of Buffer A supplemented with 1 mM β-DDM, 200 µM TCEP, 100 µM PMSF, and 50 mM Imidazole. The protein was eluted with 2.5 resin volumes of the same buffer but containing 300 mM Imidazole. Then, the buffer was exchanged in 50-kDa MWCO centrifuge concentrators to 20 mM Tris/HCl pH 7.4, 150 mM NaCl, 100 µM TCEP, 1 mM β-DDM (Buffer B), and 15 mM Imidazole, and the protein solution was mixed with Co resin (His-select Cobalt Affinity Gel, *Sigma Aldrich*) for 1h at 4 °C. The unbound fraction was discarded, and the resin was washed with 3 volumes of Buffer B containing 20 mM Imidazole. The apoAI-EfpA protein was eluted with Buffer B containing 300 mM Imidazole. Then, Imidazole was removed from the protein solution by extensive washing with Buffer B in 50-kDa MWCO centrifuge concentrators. The Imidazole-free protein solution was concentrated to ca. 25-50 µM apoAI-EfpA, frozen, and stored at -80 °C.

Size exclusion chromatography (SEC) was performed on the double affinity (Ni-+Co-affinity) purified *Mtb* EfpA using a Superdex™ 200 increase 10/300 GL size-exclusion column (*GE Healthcare*) plugged into an AKTÄ explorer 100 (*Amersham Biosciences*) protein purifier system. The buffer used was 20 mM Tris/HCl pH 7.4, 150 mM NaCl, 1 mM DDM, 200 µM TCEP, 100 µM PMSF, and 1 mM EDTA.

### Sodium dodecyl sulfate polyacrylamide gel electrophoresis (SDS-PAGE) and western blotting (WB) to detect the expression and purity of apoAI-EfpA

All *E. coli* cell fractions (insoluble fraction with inclusion bodies, membranes, and soluble fractions) were tested for the expression and localization of apoAI-EfpA using SDS-PAGE and WB. The purity of apoAI-EfpA after Ni-affinity and Co-affinity chromatography, and SEC was assessed using SDS-PAGE and WB. All the samples for SDS-PAGE and WB constituted protein, DTT (dithiothreitol) at 10 mM, and a loading buffer. The samples and marker (Precision plus protein dual-color standard, *BioRad*) were loaded onto 4%-20% Criterion™ TGX™ precast gels (*BioRad*) immersed in Tris/glycine/SDS buffer (*BioRad*), and then electrophoresis was conducted in a Midi Criterion™ vertical electrophoresis cell (*BioRad*) at 170 V. For blotting, the protein was transferred from the gel onto a 0.2-μm nitrocellulose membrane (*BioRad*) using a Criterion™ blotter (*BioRad*) overnight at 10 V and 4 °C. Protein bands on the gel were visualized by Coomassie Blue dye staining and washing with de-stain solution. We utilized colorimetric detection to visualize the apoAI-EfpA band(s) on the membrane after WB transfer: The primary mouse antihistidine tag antibody (*BioRad*) and secondary goat antimouse IgG antibody conjugated to Alkaline Phosphatase (*BioRad*) were used.

### Preparation of lipid stock solution and reconstitution of apoAI-EfpA in lipid nanoparticles

DOPC (1,2-dioleoyl-sn-glycero-3-phosphocholine; 18:1 [Δ9-Cis] PC) and DOPS (1,2-dioleoyl-sn-glycero-3-phospho-L-serine) lipids (purchased from *Avanti Polar Lipids*) in chloroform or chloroform/methanol/water were mixed in a glass vial at molar ratio of 70:30, and the solvent was evaporated under a stream of nitrogen gas. The vial was flashed until the liquid was visibly evaporated and then flashed for at least 2 h more to remove the residual organic solvent. Then a buffer containing 20 mM Tris/HCl pH 7.4 and 150 mM NaCl was added in the vial to the dry lipid film to obtain a final total lipid concentration of 30 mM. The lipid/buffer mixture was incubated for 30 min on ice to ensure complete lipid hydration. This stock solution was then used to reconstitute the apoAI-EfpA protein in lipid bilayers.

Thereafter, an aliquot of lipid stock solution was placed in a 1.5 ml tube, and the lipid was solubilized using 10% of Triton X-100 detergent in H_2_O. Then, the necessary volumes of apoAI-EfpA in β-DDM and buffer (20 mM Tris/HCl pH 7.4 and 150 mM NaCl) were added to the lipid solution to obtain a final protein-to-lipid molar ratio of 1:120 (lipid concentration of 1.8 mM, and protein concentration of 15 µM). The mixture was incubated for 1h at 22 °C. Then, BioBeads (*BioRad*) were added, and the mixture was incubated under constant mixing for 2 h at 22 °C. The BioBeads were changed 2 more times after incubation for 2 h and then overnight at 4 °C to remove the detergents completely. This proteolipid sample was used for negative-staining trasmission electron microscopy (nsEM) assesments.

### Preparation of samples for nsEM and nsEM imaging of the apoAI-EfpA protein

Prior to the negative staining, aliquots of the apoAI-EfpA in β-DDM and in lipid were placed in 1.5 ml tubes and then mixed with 5 nm Gold Nanoparticles (5-nm Gold-Ni-NTA, *Nanoprobes*) (GNPs) for 30-40 min at 4 °C. These GNPs bind to the protein His-tag.^28^ The protein-to-GNPs molar ratios were ca. 1:130 and 1:100 for protein in detergent and in lipid, respectively. We prepared and used samples of Ni+Co-affinity purified protein: GNPs-labeled protein in β-DDM (18 µM protein), nonlabeled protein in β-DDM (18 µM protein), GNPs-labeled protein in lipid (11.25 µM protein), nonlabeled protein in lipid (11.25 µM protein), and nonlabeled protein in lipid (15 µM protein); and 2 samples of SEC-purified apoAI-EfpA F1 at similar concentration and protein-to-lipid molar ratio 1:120. A 15-µl sample was loaded onto a carbon-coated copper grid and allowed to settle for 1min 30sec at room temperature. Then, the solution on the grid was removed gently using filter paper, and the settled on the grid protein in detergent or lipid was stained by adding 10 µL of 1.5% uranyl acetate (UA), incubated for 1min30sec, and the remaining UA solution was again cleared gently using filter paper. Stained samples were air dried overnight and then used for EM imaging.

The digital micrographs of negatively stained samples were collected using a transmission electron microscope (TEM *Hitachi H-7650*) equipped with a fluorescent screen with a visual field diameter of 160 mm and operated at an accelerating voltage of 60 kV with a 10-μA emission current. The direct magnification used was ×40,000-70,000. Micrographs devoid of drift and astigmatism were scanned at 1,024×1,024 pixels.

### Bioinformatic analysis of *Mtb* EfpA

The *Protter - visualize proteoform*^34^ was used to predict and visualize the *Mtb* EfpA topology in the membrane bilayer. The NCBI BLAST algorithm and the T-COFFEE Multiple Sequence Alignment software were used to predict the identities and align the amino acid sequences between EfpA exporters from various *mycobacterial* species as well as between *Mtb* EfpA and other QacA subfamily trasporters. Next, the AlphaFold2 multimer^41^ software was used to test the propensity of EfpA to self-oligomerizes, and the SWISS-MODEL software^50^ was used to generate models of *Mtb* EfpA in outward- and inward-facing conformations, which were based on homology with existing X-ray structures of NorC (PDB#7D5Q) and DtpA (PDB#6GS1).^36, 37^

## AUTHOR CONTRIBUTION

OI – experiment, figures, data analysis; AO – experiment, figures, data analysis; MMI – experiment; EH – experiment, SM – experiment, figures; OA – experiment; ERG – conception, design, experiment, data analysis and interpretation, writing the manuscript, funds acquisition, supervision. All authors contributed to the finalization of the manuscript. All authors read and approved the final version of the manuscript.

## ACKNOWLEDGMENTS

nsEM was conducted at the Art&Sciences Imaging Facility, TTU. This study was supported by a Royal Society of Chemistry Research Grant R22-2811992318 (to ERG) and start-up funds from the Department of Chemistry and Biochemistry at TTU (to ERG)

## DATA AVAILABILITY

The generated new data will be made available upon reasonable request to the corresponding author Elka R. Georgieva.

## CONFLICT OF INTERESTS

The authors declare no conflict of interests.

## Notes

### Competing Interest Statement

The authors have declared no competing interest.

### Summary of Updates

New experimental data as well as protein modeling are provided.

