## Supplementary Information for "Protein engineering, production, reconstitution in lipid nanoparticles, and initial characterization of the *Mycobacterium tuberculosis* EfpA drug exporter"

### Expression in *E. coli* inclusion bodies and refolding of *Mtb* EfpA

We designed two fusion constructs of *Mtb* EfpA that are a $His_8$-linker-FLAG-linker-*Mtb* EfpA, and $His_8$-linker-*Mtb* EfpA. The DNAs encoding these constructs were commercially synthesized and cloned in pET15b vector at NcoI and BamHI sites (GenScript, Inc), and then transformed into Bl21(DE3) *E. coli* cells. The protein was expressed using procedure similar to that described in the Main Text for the expression of apoAI-EfpA. After harvesting the *E. coli* cells and breaking them open, we screened using SDS-PAGE and western blotting (WB) each *E. coli* fraction (cytoplasmic, membrane and inclusion bodies) and found the protein in the insoluble inclusion bodies. Then we dissolved the inclusion bodies in buffer containing 20 mM-30 mM SDS [1] at room temperature (RT) for about 2 h or the mixture was heated to 27-30 °C for about 30 min. Thereafter, we added n-Dodecyl-beta-Maltoside Detergent (β-DDM) detergent to 10 mM, incubated the mixture for 30 min at RT and then placed the mixture on ice for 2 h to precipitate SDS, thus removing it from the medium and allowing for the protein to refold. Then the precipitated SDS was removed using ultracentrifugation at 30,000 rpm for 30 min in an Optima XE-90 ultracentrifuge (Beckman Coulter) in a rotor type 70.1Ti at 4 °C. The supernatant containing EfpA protein was collected, lyso lipid (mixture of 14:0 lyso PC/lyso PG) was added to 2 mM and the remaining SDS was removed by the addition of 400 mM KCl—the salt was added slowly under constant mixing. The precipitated SDS was removed again using ultracentrifugation under the same conditions. Then, His-tagged EfpA was purified using consecutively Ni- and Co-affinity purification methods. The final protein was purified to a significant extent (Figure S1).

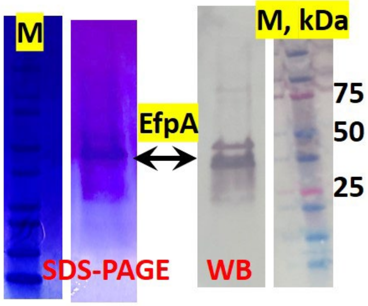


**Supporting Figure 1.** SDS-PAGE (left) and WB (right) of purified from inclusion bodies EfpA.

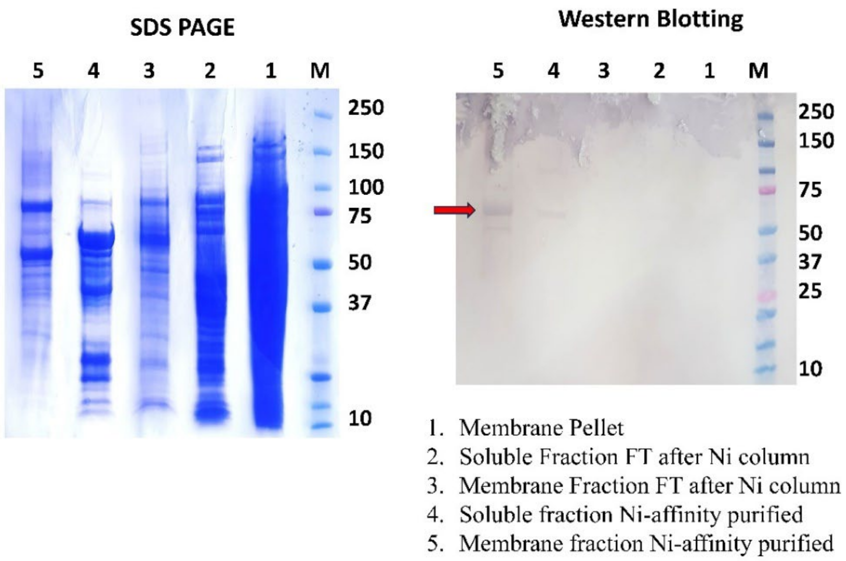


**Supporting Figure 2.** SDS-PAGE (left) and WB (right) of the *E. coli* soluble and membrane fractions after the expression of apoAI-EfpA. The protein was found mostly in the membrane fraction and to much lesser extend in the soluble fraction. The molecular weights of proteins in protein marker (M) are shown on the right.

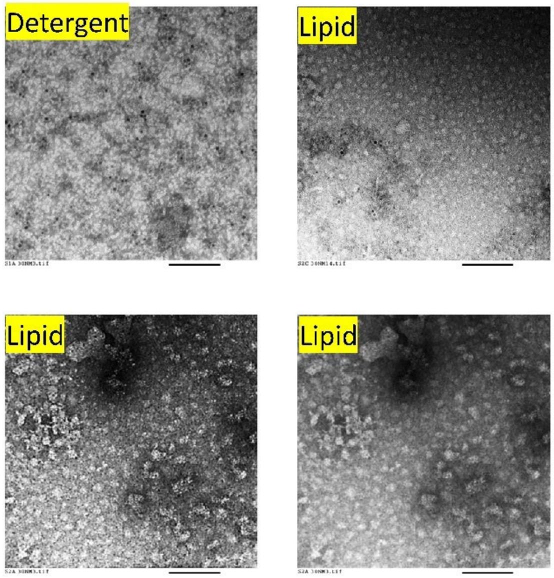


**Supporting Figure 3.** nsEM images of purified apoAI-EfpA in detergent (β-DDM) and in lipid (DOPC/DOPS). Protein particles are clearly visible in both the detergent and lipid. In lipid along with mostly discoidal protein-lipid nanoparticles, of larger protein-lipid objects (a very small fraction) are also visible.

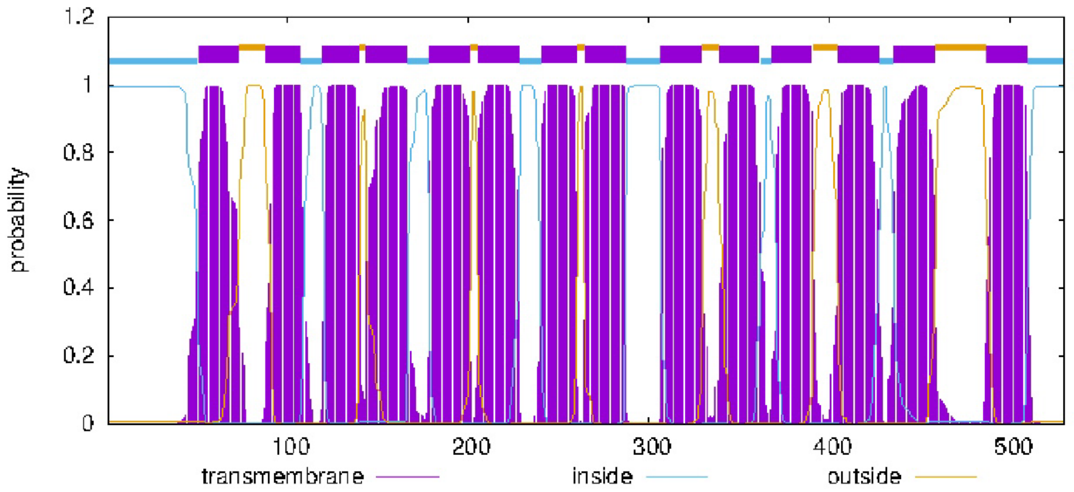


**Supporting Figure 4.** *Mtb* EfpA exporter's membrane topology was predicted using the TMHMM software. EfpA has 14-TM helices with N- and C-termini located in the intracellular space. The protein regions located inside and outside the cell are colored in blue and orange, respectively.

1. Michaux C, Pomroy NC, Prive GG. Refolding SDS-denatured proteins by the addition of amphipathic cosolvents. J Mol Biol. 2008;375(5):1477-88. Epub 20071119. doi: 10.1016/j.jmb.2007.11.026. PubMed PMID: 18083190.
